# Light tunes a novel long-term threat avoidance behavior

**DOI:** 10.1101/2024.10.28.620706

**Authors:** Marcos L. Aranda, Eric Min, Lucy T. Liu, Anika E. Schipma, Tiffany M. Schmidt

## Abstract

Animals must constantly scan their environment for imminent threats to their safety. However, they must also integrate their past experiences across long timescales to assess the potential recurrence of new threats. Though visual inputs are critical for the detection of environmental danger, whether and how visual information shapes an animal’s assessment of whether a new threat is likely to reappear in a given context is unknown. Using a novel behavioral assessment of long-term threat avoidance behavior, we find that animals will avoid a familiar location where they previously experienced a single exposure to an innately threatening visual stimulus. This avoidance behavior is highly sensitive and lasts for multiple days. Intriguingly, we find that the melanopsin-expressing, intrinsically photosensitive retinal ganglion cells tune this behavior via a perihabenular-nucleus accumbens circuit distinct from the canonical visual threat detection circuits. These findings define a specific retinal cell type driving a new long-term threat avoidance behavior driven by prior visual experience.

## Introduction

Detecting existing threats is key to animal survival, and organisms have evolved many mechanisms for threat detection and escape. However, animals entering each new or familiar environment must also constantly draw on past experience to calibrate their expectation that a new threat will appear, and then they must modify their behavior accordingly. The ability to balance risk-taking versus avoidance when making decisions based on expectation of encountering a threat has a clear evolutionary advantage for optimizing animal survival, foraging, and reproduction. Chronic over-avoidance of perceived threats in innocuous situations could result in excessive vigilance, leading to failure to forage or reproduce, while under-avoidance of perceived threats in dangerous environments could result in excessive risk-taking, leading to injury or death. The circuits underlying these avoidance calculations must therefore be plastic, well-calibrated, and operate in varied contexts and timescales. Despite their importance for survival and decision making, these threat avoidance circuits are not well understood.

Here, we introduce a behavioral paradigm to characterize and quantify avoidance of threats over long timescales based on past experience, a behavior we have termed long term threat avoidance (LTTA). We find that animals exposed just once to an innately threatening visual stimulus will then readily avoid that location days later, even in the absence of any current threat. Intriguingly, this behavior is tuned by visual circuits distinct from those involved in threat detection and requires input from the melanopsin-expressing, intrinsically photosensitive retinal ganglion cells (ipRGCs) to a retino-thalamic-limbic circuit, defining a novel circuit key for shaping avoidance behavior based on past experience.

## Results

### A paradigm to measure long-term threat avoidance in mice

We sought to measure long-term threat avoidance (LTTA) behavior by developing a novel paradigm to test the impact of a single visual looming threat exposure in a previously neutral context on an animal’s future avoidance behavior, days later, in that same context (Fig. 1A). The visual looming stimulus has many features that make it ideal to probe the circuits and mechanisms underlying threat avoidance over long timescales because 1) a single exposure to a looming stimulus drives an innate fear response through well-characterized visual circuits, 2) it is highly ethologically relevant for the mouse, and 3) it does not rely on physically aversive stimuli, which make it difficult to distinguish visual versus nociceptive contributions (Gross and Canteras, 2012; LeDoux, 2000; Tovote et al., 2015; Yilmaz and Meister, 2013). In this LTTA paradigm, dark adapted mice are placed in a square arena with a monitor overhead projecting a neutral gray background (see methods). We track each animal’s exploratory behavior during this *Pre-Exposure* phase before any stimulus is presented. During *Pre-Exposure*, we find that control animals readily explore the edges (80% of total time) and center (20% of total time) of the arena (Fig. S1). Four minutes later, animals enter the *Exposure* phase, where they are exposed to a single looming stimulus in the center of the arena, which we refer to as the ‘threat zone’ (TZ) (Evans et al., 2018; Fratzl et al., 2021). Animals then remain in the arena for 1-minute following *Exposure*, after which they are returned to their home cage. Two days later, we assess whether the prior exposure to that single looming stimulus drives avoidance of the TZ by placing animals back in the same arena for a 4-minute *Test* phase where they are allowed to freely explore the arena, but no threatening stimulus is presented (Fig. 1A).

**Figure 1.**
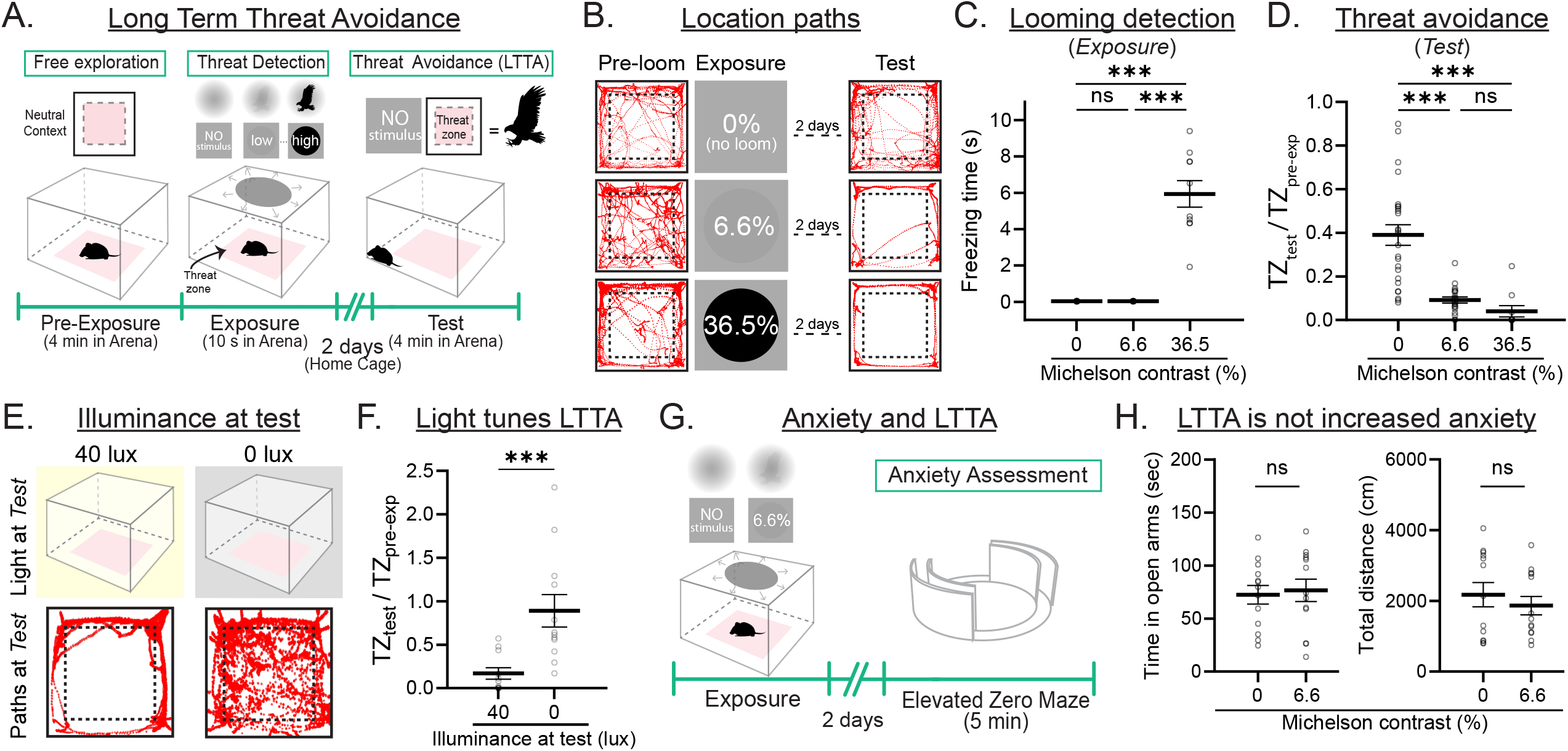
A paradigm to measure long-term threat avoidance (LTTA) in male and female mice. (A) Schematic representation of LTTA paradigm. (B) Representative location paths of control male and female mice during the *Pre-exposure* (Pre-loom), *Exposure* and *Test* phases of the LTTA paradigm at 0, 6.6 and 36.5% Michelson contrasts. (C) Freezing time at 0, 6.6 and 36.5% Michelson contrast during the *Exposure* phase of the LTTA paradigm in control male and female mice (n = 5-17 mice/group). (D) Time in threat zone (TZ) during the *Test* phase normalized to time in TZ during the *Pre-exposure* phase in control male and female mice (n = 5-17 mice/group). (E) Behavioral paradigm to test the role of light on LTTA (*top*) and representative animal position traces during the Test phase of the LTTA paradigm at 0 and 40 lux of control male and female mice (*bottom*). (F) Time in TZ during the *Test* phase normalized to time in TZ during the *Pre-exposure* phase (n = 11-12 mice/group) in control male and female mice. (G) Behavioral paradigm to test anxiety in looming-exposed mice. (H) Time in open arms (left) and total distance (right) in the Elevated Zero Maze (EZM) assay of control male and female mice previously exposed to 0 or 6.6% looming stimulus (n = 13 mice/group). All data are Mean ± SEM, n.s. (not significant) P>0.05; ***P < 0.001. Student t- and one-way ANOVA with Tukey’s multiple comparisons test.

We first analyzed looming detection of expanding disks varying in contrast from low (6.6% Michelson contrast) to high (36.5%, maximum achievable contrast for a black disk on gray, see Methods) in male and female mice by quantifying freezing behavior during the *Exposure* phase (Fig. 1B-C and S1). Both male and female mice showed detectable freezing behavior starting at 13.1% *Exposure* that increased with higher contrast. Animals showed no detectable freezing behavior at 6.6% *Exposure* (Fig. 1B-C and S1). These results defined a subthreshold contrast (6.6%) for freezing behavior and support previous findings that increased looming stimulus contrast enhances saliency to drive increased freezing behavior (Calanni et al., 2024; Evans et al., 2018; Fratzl et al., 2021).

We then tested long-term avoidance behavior by quantifying and comparing time in the TZ during the *Test* versus *Pre-Exposure* phases (Fig. 1A and see methods). Animals previously exposed to a looming stimulus of any contrast, even 6.6%, spent significantly less time in the TZ compared to 0% *Exposure* controls (Fig. 1D, S1), indicating that TZ avoidance is not simply due to habituation. We were surprised to observe robust TZ avoidance during the *Test* phase 2 days following 6.6% *Exposure* because freezing behavior was not detected during the 6.6% looming *Exposure* phase (Fig. 1B-D, S1)., These findings indicate that the circuits underlying LTTA are more sensitive and likely distinct from the well-characterized, canonical, looming detection circuits (Fratzl et al., 2021; Salay and Huberman, 2021; Shang et al., 2018; Wang et al., 2021; Wei et al., 2015). Importantly, LTTA behavior requires visual input because animals placed in the dark during the *Test* phase following a 6.6% *Exposure* fail to avoid the threat zone, in stark contrast to animals placed in standard 40 lux illumination during the *Test* phase following an identical, 6.6% *Exposure* (Fig. 1E-F, S2). Moreover, LTTA is not due to increased general anxiety because mice showed no changes in behavior in the Elevated Zero Maze 2 days post 6.6% *Exposure* (Fig. 1G-H, S3). Thus, the circuits underlying long-term threat avoidance are highly sensitive, long-lasting, and distinct from those driving anxiety and looming-dependent freezing responses.

Notably, male and female mice show robust TZ avoidance in the LTTA assay that requires visual input and is independent of general anxiety levels (Fig. S2, S3). However, when we probed female LTTA across stages of estrus, we noted extensive variation in avoidance behavior in sexually receptive (SR) versus non-sexually receptive (NR) stages (Fig. S1), suggesting interplay between reproductive drive and exploratory behavior in females. For this reason, we chose to continue our initial circuit characterization only in male mice.

### ipRGCs tune long-term threat avoidance

We next sought to identify the retinal cell types involved in LTTA. Given that LTTA is highly dependent on environmental luminance (Fig. 1E-F, S2), the melanopsin-(Opn4)-expressing intrinsically photosensitive retinal ganglion cells (ipRGCs) are a good candidate for tuning LTTA because melanopsin signaling within ipRGCs encodes environmental luminance over a large dynamic range and across long timescales (Aranda and Schmidt, 2020; Berson et al., 2002; Hattar et al., 2002). To determine whether melanopsin signaling is required for LTTA, we compared TZ avoidance of melanopsin null (Opn4^-/-^) animals to littermate control animals following 6.6% or 0% *Exposure*. 6.6% looming contrast was chosen to isolate impacts on avoidance behaviors versus detection. Indeed, Opn4^-/-^ and control littermates show no freezing behavior at either contrast (Fig. S4). Following 6.6% *Exposure*, Opn4^-/-^ animals spent significantly more time in TZ compared to controls, indicating that melanopsin signaling in ipRGCs is required for LTTA (Fig. 2A-C, S4). Importantly, Opn4^-/-^ mice at 0% *Exposure* showed similar time in the TZ to control littermates in both the *Pre-exposure* and *Test* phases (Fig. S4), indicating that differences in LTTA are not due to differences in baseline exploratory behavior (Fig. 2B-C, S4). Additionally, these differences are not due to altered anxiety because Opn4^-/-^ mice showed no differences in anxiety-like behaviors compared to control littermates (Fig. S5).

**Figure 2.**
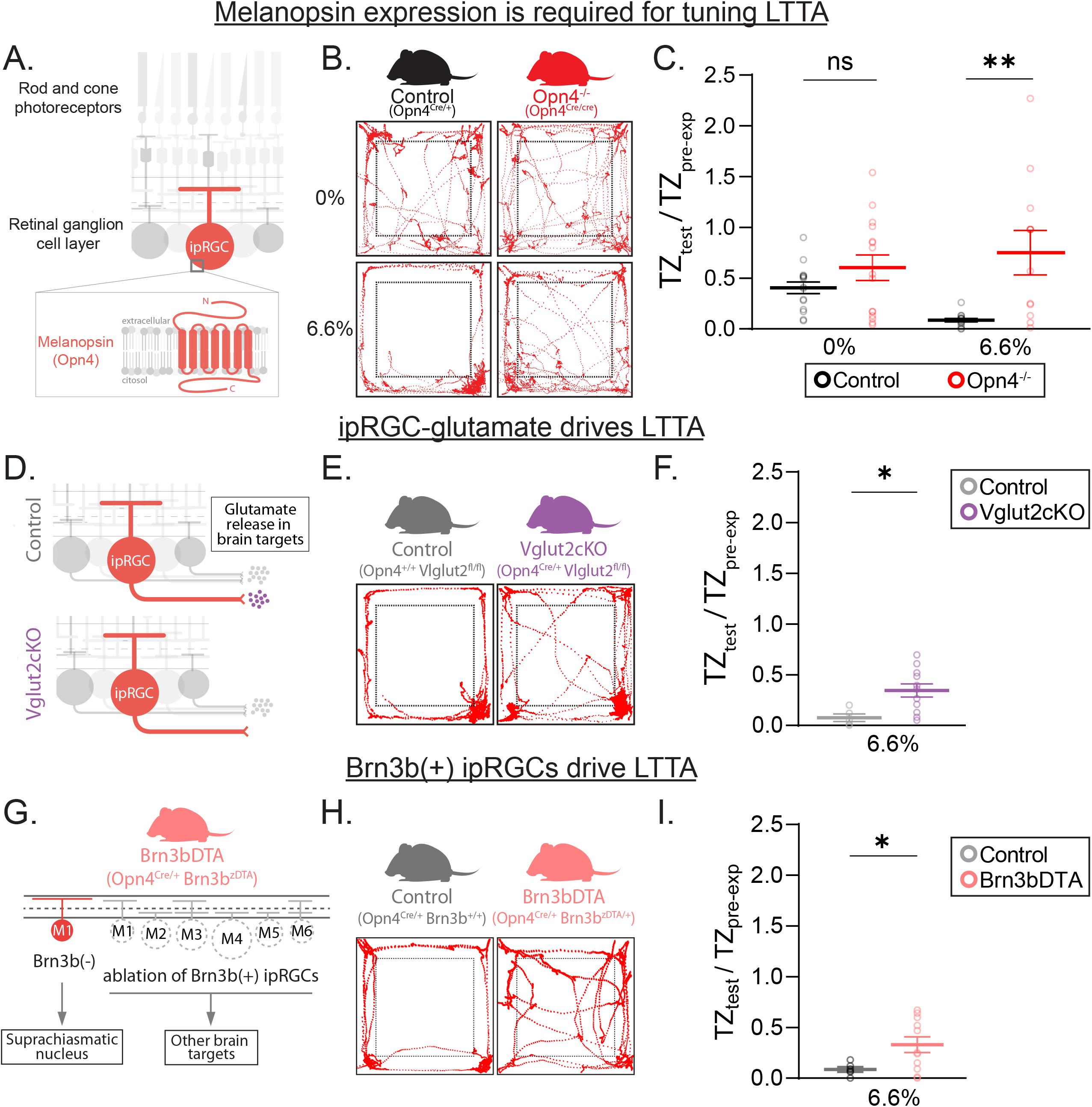
ipRGCs drive LTTA. (A) Schematic representations of mammalian retina and specific melanopsin expression in ipRGCs. (B) Representative animal position traces of control and melanopsin-null (Opn4^-/-^) male mice during the *Test* phase of the LTTA previously exposed to 0 or 6.6% Michelson contrast. (C) Time in threat zone (TZ) during the *Test* phase normalized to time in TZ during the *Pre-exposure* phase in control and Opn4^-/-^ male mice exposed to 0 or 6.6% Michelson contrast (n = 11-15 mice/group). (D) Schematic representations of glutamate release ablation specifically from ipRGCs. (E) Representative animal position traces of Vglut2cKO male mice during the *Test* phase of the LTTA previously exposed to 6.6% Michelson contrast. (F) Time in TZ during the *Test* phase normalized to time in TZ during the *Pre-exposure* phase in Vglut2cKO and control male mice exposed to 6.6% Michelson contrast (n = 11-15 mice/group). (G) Schematic representations of Brn3b-positive ipRGC ablation in Brn3bDTA mice. (H) Representative animal position traces during the *Test* phase of the LTTA paradigm. (I) Time in TZ during the *Test* phase normalized to time in TZ during the *Pre-exposure* phase of Brn3bDTA and control littermate male mice (n = 5-11 mice/group). All data are Mean ± SEM, n.s. (not significant) P>0.05; *P < 0.05; **P < 0.01. Student t- and Mann-Whitney U tests.

If ipRGCs tune LTTA then disrupting ipRGC glutamate signaling should alter LTTA. Indeed, conditional knockout of the vesicular glutamate transporter Vglut2 in ipRGCs (Vglut2cKO: Opn4^Cre/+^; Vglut2^fl/fl^) caused similar deficits in TZ avoidance to Opn4^-/-^ animals despite similar baseline exploratory behavior (Keenan et al., 2016) (Fig. 2D-F, S6). Moreover, ablation of non-SCN-projecting Brn3b-positive ipRGCs using the Brn3bDTA line (Opn4^Cre/+^; Brn3bz^DTA/+^) (Chen et al., 2011), resulted in similar deficits (Fig. 2G-I, S6). Combined, these results indicate that ipRGCs release glutamate at non-SCN targets to tune LTTA behavior.

### The perihabenular nucleus is required for tuning LTTA

We next sought to identify ipRGC-recipient targets required for LTTA using an unbiased cFos screen of brain regions differentially activated during the *Test* phase in control and Opn4^-/-^ animals (Fig. 3A). If a given ipRGC-recipient region is required for LTTA, then we would expect that target to show differences in cFos activation that correlate with those seen in behavior (i.e. cFos levels in all groups are similar with the exception of 6.6% *Exposed* controls). Remarkably, the only ipRGC-recipient target showing the predicted differences in cFos labeling was the perihabenular nucleus (PHb) (Fig. 3B-C), suggesting an ipRGC-PHb circuit may underly LTTA. Notably, regions implicated in the canonical looming circuit such as medial superior colliculus (mSC) and the ventral lateral geniculate nucleus (vLGN) (Fratzl et al., 2021; Li et al., 2023; Salay and Huberman, 2021) and ipRGC-recipient regions that directly connect to looming detection circuits showed no differences in cFos expression following LTTA (Fig. S7), again suggesting that the LTTA circuit architecture is distinct from the looming detection circuit.

**Figure 3.**
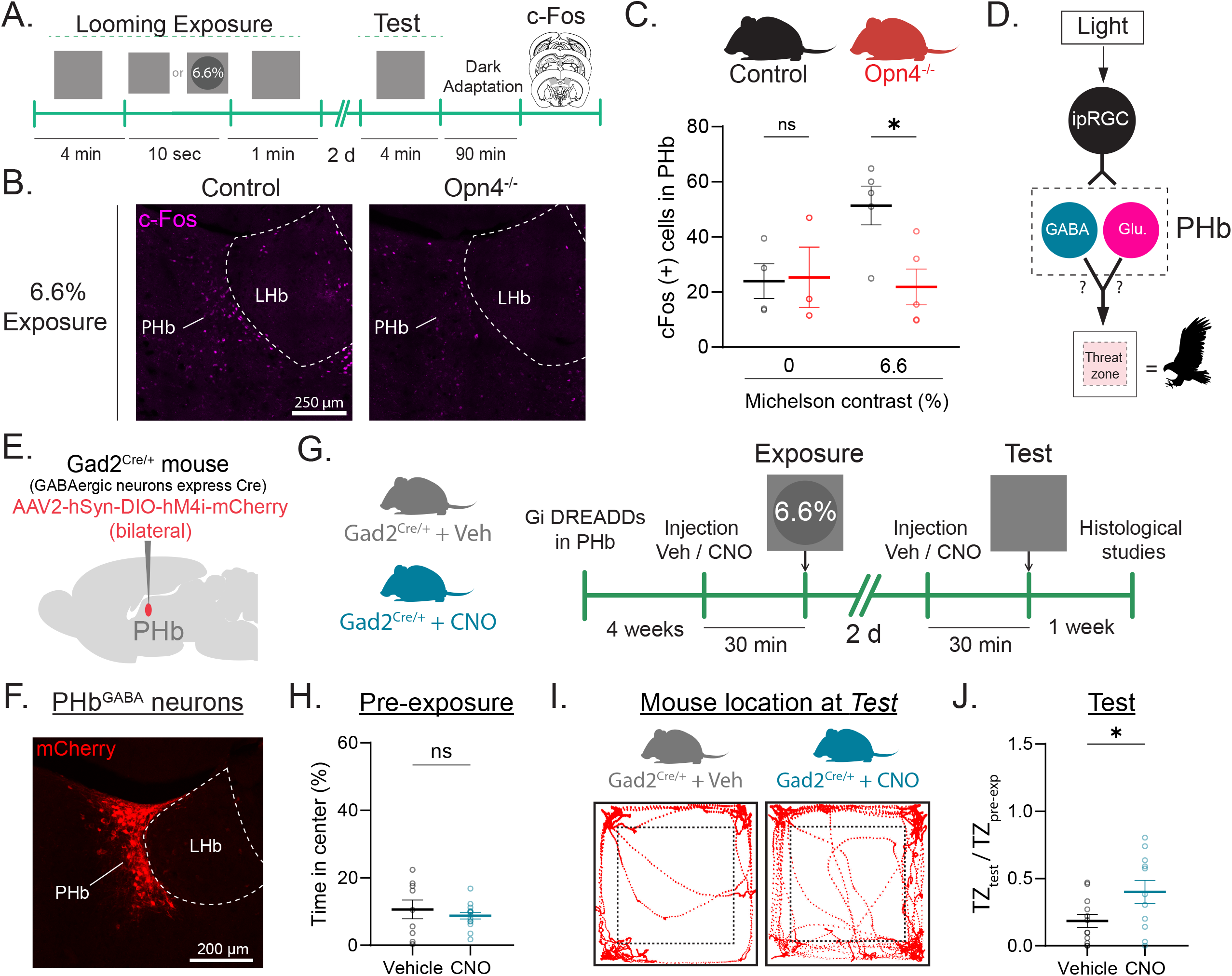
PHb^GABA^ are required for LTTA. (A) Schematic representation of experimental design for cFos staining. (B) Representative images of cFos staining in PHb from control and melanopsin-null (Opn4^*-/-*^) male mice exposed to 6.6% Michelson contrast. (C) cFos-positive cells quantification in the PHb from control and control and Opn4^-/-^ male mice exposed to 0 and 6.6% Michelson contrast (n = 3-5 mice/group). (D) Schematic representation of potential ipRGC-PHb circuit driving LTTA. (E) Scheme of viral injection of Gi-DREADDs in PHb of Gad2^Cre^ male mice. (F) Representative image of Gi-DREADDs expression in PHb. (G) Time of experimental design. (H) Percentage of time in center during the Pre-exposure phase in Vehicle or CNO-injected Gad2^Cre^ male mice (n = 9-14 mice/group). (I) Representative Gad2^Cre^ animal position traces during the *Test* phase of the LTTA paradigm. (J) Time in threat zone (TZ) during the *Test* phase normalized to time in TZ during the *Pre-exposure* phase of Vehicle and CNO-injected Gad2^Cre^ male mice (n = 11-12 mice/group). All data are Mean ± SEM, n.s. (not significant) P>0.05; *P < 0.05. Student t- and Mann-Whitney U tests.

ipRGCs primarily innervate GABAergic neurons in the caudal PHb (PHb^GABA^), which in turn make local connections with excitatory-relay PHb neurons (PHb^Glu^) and anterior PHb^GABA^ neurons (Fig. 3D) (An et al., 2020; Fernandez et al., 2018; Weil et al., 2022). If PHb^Glu^ and/or PHb^GABA^ neurons are required for LTTA, then silencing that specific population should disrupt LTTA. To test this, we injected Cre-dependent Gi-DREADDs in the PHb of Gad2^Cre^ and Vglut2^Cre^ mice to silence either PHb^GABA^ (Fig. 3E-F, see Table S1) or PHb^Glu^ (Fig. S8) neurons, respectively. We then intraperitoneally injected either vehicle or the DREADD agonist clozapine-N-oxide (CNO) 30 minutes prior to both the *Pre-Exposure* and *Test* phases of the LTTA paradigm (Fig. 3G). Importantly, CNO injection did not alter exploratory behavior during *Pre-Exposure* phase compared to vehicle-injected mice, indicating that CNO itself does not alter baseline exploratory behavior (Fig. 3H). Silencing of PHb^GABA^ (Fig. 3E-J), but not PHb^Glu^ neurons (Fig. S8) disrupted LTTA, causing animals to spend significantly more time in the TZ, confirming that the GABAergic neurons of the ipRGC-recipient PHb are required for LTTA.

If PHb-ipRGCs are required for LTTA, then activation of PHb-ipRGCs in melanopsin null animals should rescue the observed deficits in LTTA in Opn4^-/-^ animals. We have previously shown that Gq activation using Gq-DREADDs in ipRGCs *ex vivo* restores melanopsin-like photoresponses in melanopsin null ipRGCs (Sonoda et al., 2018). We therefore adapted this approach to our *in vivo* LTTA paradigm by bilaterally injecting retrograde (rg)AAV into the PHb of melanopsin null (Opn4^Cre/Cre^) animals to drive Cre-dependent expression of Gq-DREADDs (rgAAV-hSyn-DIO-hM3Gq-mCherry) selectively in PHb-ipRGCs (Fig. 4A, see Table S1). We then intraperitoneally injected either vehicle or CNO 30 minutes prior to both the *Pre-Exposur*e and *Test* phases (Fig. 4A). CNO-injected Opn4^-/-^ animals avoided the TZ, spending significantly less time in the TZ than vehicle-injected melanopsin null littermates. This indicates that Gq-DREADD activation of PHb-ipRGCs restores LTTA (Fig. 4B-D). CNO administration did not alter baseline exploratory behavior during the *Pre-Exposure* phase compared to vehicle-injected mice (Fig. S9), again suggesting that CNO does not affect baseline exploratory behavior. These effects are not through collateral outputs of PHb-ipRGCs to the ventral lateral geniculate nucleus (vLGN), a collateral target of PHb-ipRGCs (Fig. S10, see Table S1) that has been implicated in looming-response behavior (Fratzl et al., 2021; Salay and Huberman, 2021), because chemogenetic silencing of the vLGN itself did not affect LTTA in WT mice (Fig. S11, see Table S1). Collectively, these findings indicate that PHb-ipRGCs are required for LTTA.

**Figure 4.**
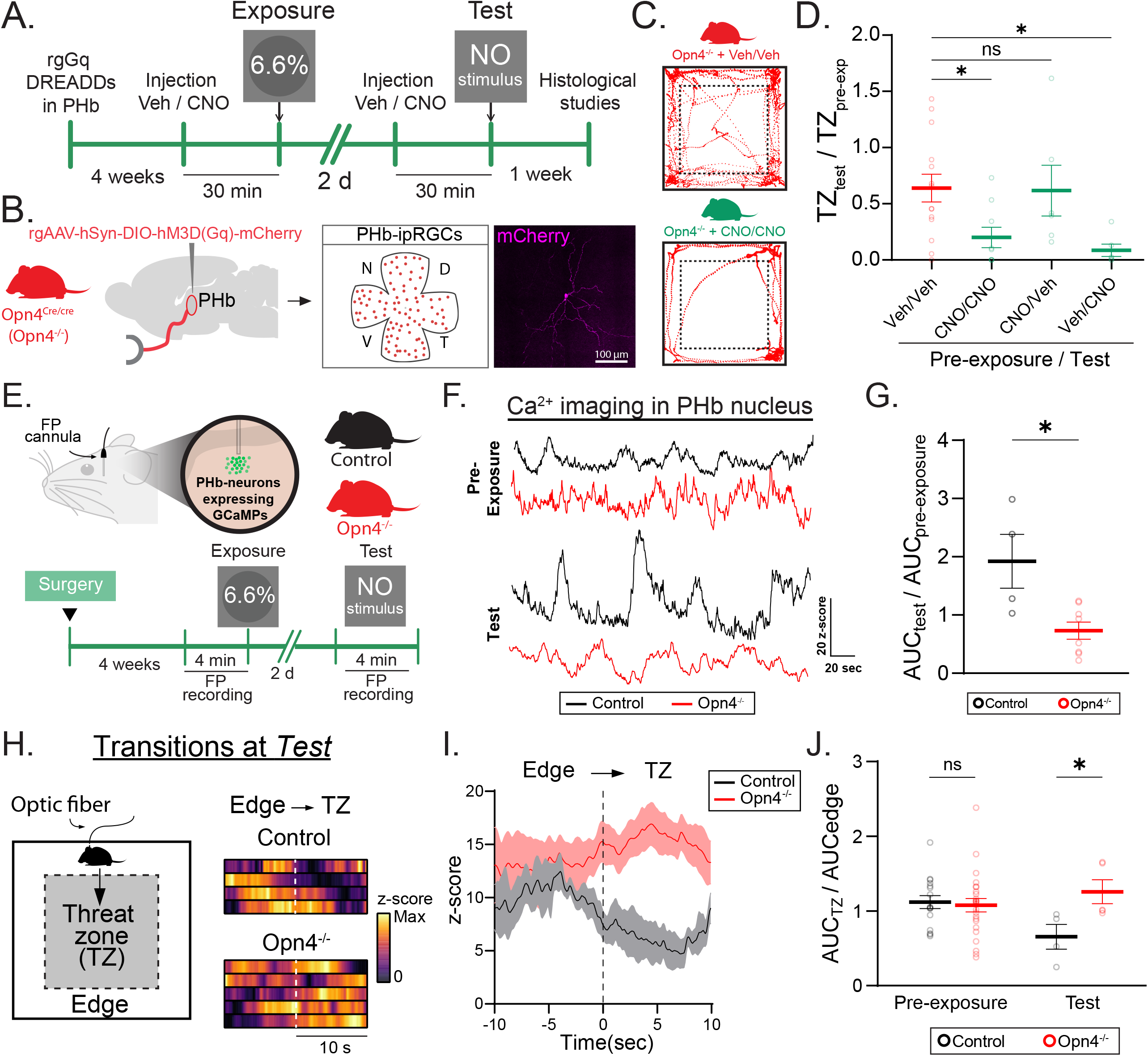
ipRGCs tune PHb neuron activity during re-exposure to context following threat exposure. (A) Scheme of viral injection of rgGq-DREADDs in PHb of Opn4^-/-^ male mice. (B) Representative images of Gq-DREADDs expression in a PHb-ipRGC. (C) Representative Opn4^-/-^ animal position traces during the *Test* phase of the LTTA paradigm. (D) Time in trheat zone (TZ) during the *Test* phase normalized to time in TZ during the *Pre-exposure* phase of Vehicle or CNO-injected Opn4^-/-^ male mice (n = 6-14 mice/group). (E) Scheme of experimental design for fiber photometry (FP) in LTTA paradigm. (F) Representative Ca^2+^ signal recordings from PHb in control and Opn4^-/-^ male mice during *Pre-exposure* and *Test* phases of the LTTA paradigm. (G) Area under the curve (AUC) of *z*-score of FP signal during the *Test* phase normalized to AUC of *z*-score of FP signal during the *Pre-exposure* phase in control and Opn4^-/-^ male mice (n = 4-6 mice/group). (H) Schematic representation of transitions from edge to TZ (left) and heatmaps of *z*-score of FP signal in animal transitions from edge to TZ during the *Test* phase of the LTTA paradigm in control and Opn4^-/-^ male mice (n = 4-5 mice/group) (right). (I) *z*-score of FP signal in animal transitions from edge to TZ during the *Test* phase of the LTTA paradigm in control and Opn4^-/-^ male mice (n = 4-5 mice/group). (J) Averaged AUC of *z*-score of FP signal when mice were placed in TZ normalized to AUC of z-score of FP signal when mice were placed in edges during transitions in the *Pre-exposure* and *Test* phases of the LTTA paradigm (n = 4-27 transitions/group). All data are Mean ± SEM, n.s. (not significant) P>0.05; *P < 0.05. Student t-, Mann-Whitney U tests, and one-way ANOVA with Dunn’s multiple comparisons test.

### ipRGCs tune PHb neuron activity to facilitate recall of the threat/context association

To determine whether PHb-ipRGCs are important for LTTA during the formation of association between threat and context (i.e. at *Pre-Exposure / Exposure* phases) or whether they facilitate the recall of this association upon re-exposure (i.e. at *Test* phase), we bilaterally injected retrograde (rg)AAV-Gq-DREADDs into the PHb of melanopsin null (Opn4^Cre/Cre^) animals, but administered a single dose of CNO prior to *Pre-Exposure* (CNO/Veh group) *or Test* (Veh/CNO group) phases to *Opn4*^*-/-*^ mice (Fig. 4A-D, see Table S1). A third group received two vehicle injections (Veh/Veh group) as control. Animals that received CNO prior to the *Test* phase (Veh/CNO), but not those that received CNO prior to the *Pre-Exposure* phase (CNO/Veh), showed decreased time in the TZ (Fig. 4D). These results paralleled those where animals received CNO in both the *Pre-Exposure* and *Test* phases and indicate that PHb-ipRGC activity is required for LTTA only during the *Test* phase when the animal must recall the association between threat and context.

To understand PHb signaling dynamics and the impact of melanopsin signaling during LTTA, we next used fiber photometry (FP) to measure bulk Ca^2+^ activity in the PHb *in vivo* in Opn4^-/-^ and control littermates at different phases of the LTTA paradigm. To do this, we injected an AAV containing GCamp6s (AAV1-CAG-GCaMP6s-WPRE-SV40) and implanted a FP canula unilaterally in the PHb of control and Opn4^-/-^ mice (Fig. 4E, S12, see Table S1 and methods). We then compared overall activity in the *Pre-Exposure* versus *Test* phases of the LTTA paradigm using a 6.6% contrast looming stimulus. In agreement with cFos expression analysis (Fig. 3C), we found decreased PHb activity in Opn4^-/-^ mice compared to control littermates during the *Test* phase, but not during the *Pre-exposure*, phase of the LTTA paradigm. This indicates that ipRGC signaling during the *Test* phase serves to increase baseline PHb activity (Fig. 4F-G).

We next examined how PHb neuron activity varied with animal behavior in the LTTA paradigm. To identify key parameters for further analysis, we performed a principal component analysis (PCA) of bulk Ca^2+^ activity in the PHb and behavioral parameters data (Fig. S12). We observed that the *z*-score parameter of the FP recordings was highly correlated with behavioral parameters when mice remained in the edges (i.e. increased avoidance) compared to those behavioral parameters when mice were in the TZ (Fig. S12). To better understand this, we analyze FP signal specifically while control and Opn4^-/-^ animals occasionally transitioned from the edge to the TZ during the *Test* phase (Fig. 4H). In control mice, we found a significant decrease in the Ca^2+^ signal when control mice transitioned from the edge to the TZ, but no significant change during transitions of Opn4^-/-^ mice (Fig. 4I-J). These changes were observed only during the *Test* phase (Fig. 4J, S12), suggesting again that PHb signaling is required to facilitate the recall of the threat/context association. These data indicate that the PHb nucleus is key for encoding edge-TZ transitions and that this process requires melanopsin signaling within ipRGCs.

### A PHb to nucleus accumbens (NAc) circuit is required for LTTA

The PHb nucleus is part of a thalamo-frontocortico-striatal loop projecting to the ventromedial prefrontal cortex (vmPFC) and the nucleus accumbens (NAc) (Fig. 5A-C) (An et al., 2020; Fernandez et al., 2018; Weil et al., 2022). We therefore next tested whether PHb-NAc or PHb-vmPFC neurons are required for LTTA. If PHb-NAc or PHb-vmPFC neurons are required for LTTA, then activating them with Gq-DREADDs in Opn4^-/-^ animals should rescue LTTA deficits in these mice. To test this, we expressed excitatory Gq-DREADDs in vmPFC-, or NAc-projecting PHb neurons by bilaterally injecting a rgAAV-pkg-Cre into the NAc-or vmPFC- and a Cre-dependent Gq-DREADD (AAV-hSyn-DIO-hM3Gq-mCherry, see Table S1 and methods) into the PHb of Opn4^-/-^ mice (Fig. 5D, S13). Chemogenetic manipulation of PHb-NAc circuit induced decreased time in the TZ (i.e. increased avoidance) in CNO-injected Opn4^-/-^ mice compared to vehicle-injected Opn4^-/-^ littermate controls (Fig. 5D-I). However, we did not observe differences in LTTA when PHb-vmPFC was chemogenetically excited in Opn4^-/-^ mice (Fig. S13). These results suggest that the PHb-NAc circuit, and not PHb-vmPFC, is sufficient for LTTA in mice.

**Figure 5.**
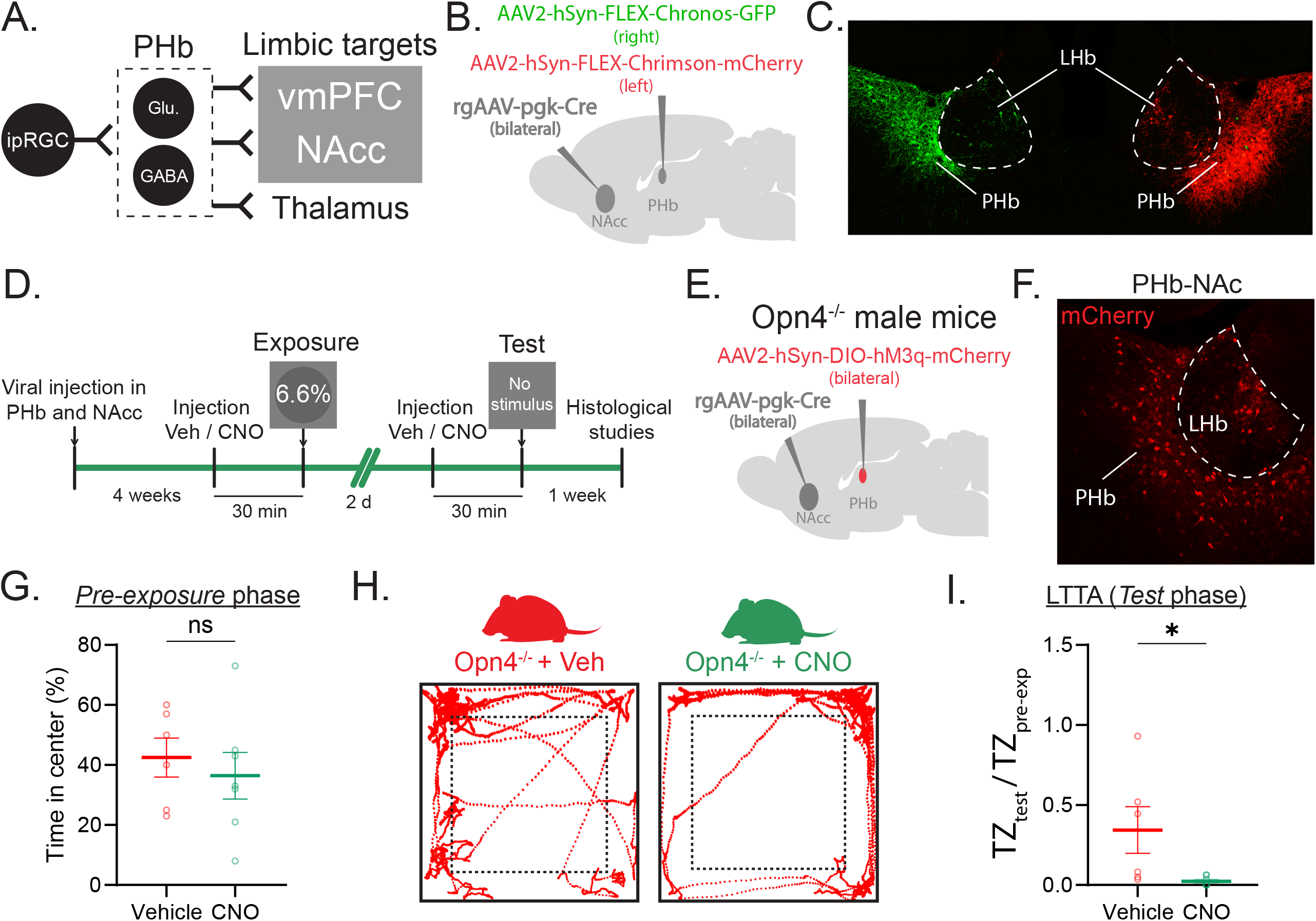
PHb-NAc downstream circuit is required for LTTA. (A) Schematic representation of PHb-downstream regions. (B) Scheme of viral injection to intersectionally trace the PHb-NAc circuit. (C) Representative image of PHb neuron axons innervating the right (green) and the left (red) NAc (n = 2 mice). (D) Scheme of experimental design. (E) Scheme of viral injection to interjectionally express Gq-DREADDs in PHb projecting to NAc in Opn4^-/-^ male mice. (F) representative image of Gq-DREADD expression in PHb-NAc neurons. (G) Time in center in vehicle and CNO-injected Opn4^-/-^ mice during the *Pre-exposure* phase of the LTTA paradigm (n = 6-7 mice/group). (H) Representative Vehicle and CNO-injected Opn4^-/-^ animal position traces during the *Test* phase of the LTTA paradigm. (I) Time in threat zone (TZ) during the *Test* phase normalized to time in TZ during the *Pre-exposure* phase of Vehicle and CNO-injected Opn4^-/-^ male mice (n = 6-7 mice/group). All data are Mean ± SEM, n.s. (not significant) P>0.05; *P < 0.05. Student t- and Mann-Whitney U tests.

## Discussion

The ability to balance and calibrate behaviors that promote risk versus safety are key to animal survival. Here we describe LTTA, a visual, ethologically relevant, long-term avoidance behavior in which animals exposed to a single, low contrast looming stimulus that was insufficient to drive a detectable looming response (i.e. freezing), caused avoidance of the TZ days following threat exposure. The circuit underlying LTTA is distinct from the looming detection or anxiety circuits and requires ipRGC projections to the recently identified perihabenular-nucleus accumbens circuit. Notably, the LTTA behavior is robust in both male and female mice. However, we observed an estrus-stage dependent impact of LTTA in females, indicating an interaction between reproductive drive and TZ avoidance behavior, supported by strong TZ avoidance in females only in the non-sexually receptive stages of estrus. These findings suggest that the LTTA circuit is a potential target of sex hormones and this intersection of ipRGCs and sex hormone modulation will be a key topic for future study.

Excitingly, we identified a previously unappreciated role of ipRGCs in facilitating the recall of the association between threat and environment by increasing baseline PHb activity during the threat avoidance behavior days after the threat exposure. Specifically, our *in vivo* recordings of PHb calcium signals during LTTA indicate that the PHb may be particularly important for encoding arena location during edge-TZ transitions, and that melanopsin signaling within ipRGCs are critical for this process. Importantly, while our results indicate that ipRGCs are dispensable to form the association between threat and environment, the retina-brain circuits necessary to encode that association during the *Pre-exposure* and *Exposure* phases of the LTTA paradigm remain to be elucidated, as does the intersection of the LTTA and looming stimulus detection circuits.

Together, our study opens new insights into the influence of retina-brain circuits on threat avoidance behaviors and paves the way for future studies on how light calibrates decision making in threatening contexts.

## Acknowledgments

National Institutes of Health grant R01 EY030565 and National Institutes of Health grant DP2 EY022584 (TMS). We thank Dr. Samer Hattar for the gift of Opn4^Cre/+^ and Dr. William Klein for the gift of Brn3bDTA mice.

## Author contributions

Conceptualization: MLA and TMS.

Methodology: MLA, EM, LL, AES.

Investigation: MLA, EM, LL, AES and TMS.

Visualization: MLA and TMS.

Funding acquisition: TMS.

Project administration: TMS.

Supervision: MLA and TMS.

Writing – original draft: MLA and TMS.

Writing – review & editing: MLA and TMS.

## Declaration of interests

The authors declare no competing interests.

## Figure legends

**Figure S1. LTTA behavior is present in male and female mice**. (A) Schematic representation of LTTA paradigm. (B) Percentage of time in center during the *Pre-exposure* phase in control male mice (n = 28). (C) Freezing time during the *Exposure* phase at 0, 6.6, 13.1, 27.5 and 36.5% Michelson contrast in control male mice (n = 5-17 mice/group). (D) Time in treat zone (TZ) during the *Test* phase normalized to time in TZ during the *Pre-exposure* phase (n = 5-17 mice/group). (E-G) Percentage of time in center during the *Pre-exposure* phase (E) Representative vaginal smears images of each estrous cycle phase. Black arrowhead: nucleated epithelial cell. Black arrow: cornified squamous epithelial cell. Red arrowhead: leukocyte. (F) Representative sexually receptive (SR) and non-sexually receptive (NR) control and Opn4^-/-^ females position traces during the *Test* phase of the LTTA paradigm after 6.6% *Exposure* (left) and time in TZ during the *Test* phase normalized to time in TZ during the *Pre-exposure* phase of estrous-tracked control female mice after 6.6% *Exposure* (n = 11-18 mice/group). All data are Mean ± SEM, ns: P>0.05, *P<0.05, **P<0.01, ***P<0.001, one-way ANOVA with Dunn’s multiple comparisons test.

**Figure S2. Light tunes LTTA in male and female mice**. (A-B) Time in TZ during the *Test* phase at 0 and 40 lux normalized to time in TZ during the *Pre-exposure* phase in male (A) and female (B) mice. All data are Mean ± SEM, *P < 0.05, **P < 0.01. Student t-test.

**Figure S3. LTTA behavior is not due to general increases in anxiety levels**. (A) Behavioral paradigm. (B-C) Time in open arms (B) and total distance (C) in the Elevated Zero Maze (EZM) assay of male and female mice previously exposed to 0 or 6.6% Michelson contrast (n = 5-7 mice/group). All data are Mean ± SEM, ns: P>0.05, Student’s t-test.

**Figure S4. ipRGCs tunes threat avoidance behavior at multiple looming contrasts**. (A) Long term Threat Avoidance (LTTA) paradigm. (B) Percentage of time in center during the *Pre-exposure* phase in control and Opn4^-/-^ male mice (n = 26-28 mice/group). (C) Freezing time during the *Exposure* phase at 0, 6.6, 13.1, 27.5 and 36.5% Michelson contrast in control and Opn4^-/-^ male mice (n = 5-17 mice/group). (D) Time in treat zone (TZ) during the *Test* phase normalized to time in TZ during the *Pre-exposure* phase (n = 5-17 mice/group). All data are Mean ± SEM, ns: P>0.05, **P<0.01, ***P<0.001, Student’s t- and Mann Whitney U tests.

**Figure S5. LTTA behavior in Opn4**^**-/-**^ **mice is not due to general increases in anxiety levels**. (A) Behavioral paradigm. (B) Time in open arms (left) and total distance (right) in the Elevated Zero Maze (EZM) assay of control and Opn4^-/-^ male mice previously exposed to 0 or 6.6% Michelson contrast (n = 5-7 mice/group). All data are Mean ± SEM, ns: P>0.05, Student’s t-test.

**Figure S6. Vlgut2cKO and Brn3bDTA mice show similar baseline exploratory behavior to their control littermates**. (A) Percentage of time in center during the *Pre-exposure* phase in control (Opn4^+/+^ Vglut2^fl/fl^) and Vglut2cKO (Opn4^Cre/+^ Vglut2^fl/fl^) littermate male mice (n = 6-11 mice/group). (B) Percentage of time in center during the *Pre-exposure* phase in control (Opn4^Cre/+^ Brn3b^+/+^) and Brn3bDTA (Opn4^Cre/+^ Brn3b^zDTA/+^) littermate male mice (n = 5-11 mice/group). All data are Mean ± SEM, ns: P>0.05. Mann Whitney U tests.

**Figure S7. cFos expression patterns after LTTA paradigm in candidate retino-recipient brain regions**. mSC: medial superior colliculus, vLGN: ventral lateral geniculated nucleus. MEA: medial amygdala. PAG: periaqueductal gray. All data are Mean ± SEM, ns: not significant, P>0.05. Student’s t- and Mann Whitney U tests.

**Figure S8. PHb-glutamatergic neurons (PHb**^**Glu**^**) are not required for LTTA. (A) Scheme of viral injection in PHb**. (B) Representative picture of Gi-DREADDs expression in PHb. (C) Scheme of experimental design. (D) Percentage of time in center during the *Pre-exposure* phase in Vehicle or CNO-injected Vglult2^Cre^ male mice (n = 7-8 mice/group). (E) Representative animal position traces during the *Test* phase of the LTTA paradigm. (F) Time in threat zone (TZ) during the *Test* phase normalized to time in TZ during the *Pre-exposure* phase of Vehicle and CNO-injected Vlut2^Cre^ male mice (n = 7-8 mice/group). All data are Mean ± SEM, ns: P>0.05, Student’s t- and Mann Whitney U tests.

**Figure S9. CNO administration does not affect baseline exploratory behavior in melanopsin-null (Opn4**^**-/-**^**) male mice**. (A) Percentage of time in center during the *Pre-exposure* phase in Vehicle or CNO-injected Opn4^Cre/cre^ male mice (n = 9-12 mice/group). (B) Representative animal position traces during the *Test* phase of the LTTA paradigm of CNO-injected mice only before *Exposure* (left) or *Test* (right) phases. All data are Mean ± SEM, ns: P>0.05, Student’s t-test.

**Figure S10. PHb-ipRGCs send collateral axons to the lateral geniculate nucleus (LGN)**. (A) Upper panel: Schematic representation of anterograde labeling of ipRGC axons. Bottom: ipRGC axons innervating the PHb. (B) Left: schematic representation of viral injections to label ipsi- and contralateral PHb-collateral axons. Right: PHb-collateral axons innervating the ventral LGN (vLGN) and sparsely labeling the dorsal LGN (dLGN, white arrowheads).

**Figure S11. vLGN is not required to drive LTTA. (A) Scheme of viral infection in vLGN of WT male mice**. (B) Representative picture of Gi-DREADDs expression in vLGN. (C) Scheme of experimental design. (D) Percentage of time in center during the *Pre-exposure* phase in Vehicle or CNO-injected WT male mice (n = 5-7 mice/group). (E) Left: Representative animal position traces during the *Test* phase of the LTTA paradigm. Right: Time in threat zone (TZ) during the *Test* phase normalized to time in TZ during the *Pre-exposure* phase of Vehicle and CNO-injected WT male mice (n = 5-7 mice/group). All data are Mean ± SEM, ns: P>0.05, Student’s t- and Mann Whitney U tests.

**Figure S12. Fiber photometry Ca**^**2+**^ **signal in PHb correlates with LTTA behavioral parameters**. (A) Representative image of GCaMPs expression and optic fiber implantation in the PHb. (B) Proportion of variance of the principal component analysis (PCA) combining behavioral and fiber photometry (FP) data. (C) PCA loadings showing behavioral and FP parameters. (D) z-score of FP signal in animal transitions from edge to TZ during the *Pre-exposure* phase of the LTTA paradigm in control and Opn4^*-/-*^ male mice (n = 18-27 mice/group). All data are Mean ± SEM.

**Figure S13. PHb-vmPFC circuit is not required to drive LTTA. (A) Scheme of intersectional viral infection of PHb-vmPFC neurons in Opn4**^**-/-**^ **male mice**. (B) Representative picture of Gq-DREADDs expression in PHb. (C) Scheme of experimental design. (D) Percentage of time in center during the *Pre-exposure* phase in Vehicle or CNO-injected Opn4^-/-^ male mice (n = 5-6 mice/group). (E) Representative animal position traces during the *Test* phase of the LTTA paradigm. (F) Time in threat zone (TZ) during the *Test* phase normalized to time in TZ during the *Pre-exposure* phase of Vehicle and CNO-injected Opn4^-/-^ male mice (n = 5-6 mice/group All data are Mean ± SEM, ns: P>0.05, Student’s t-test.

## Methods

### Animals

All procedures were approved by the Animal Care and Use Committee at Northwestern University. P60-P120 male and female littermate mice were used for this study. For circuit tracing we used WT and Opn4^Cre^ (RRID:IMSR_JAX:035925) mice. For behavior, chemogenetic and histological studies, we used *WT, Opn4*^*Cre*^, *Opn4*^*Cre/+*^; *Vglut2*^*fl/fl*^ (MGI:4879031), Brn3bDTA (RRID:IMSR_JAX:038952), *Gad2*^*Cre*^ (IMSR_JAX:028867) and *Vglut2*^*Cre*^ (RRID:IMSR_JAX:016963) male mice. For fiber photometry experiments we used *Opn4*^*Cre*^ male mice.

### Long-term threat anticipation paradigm

The experimental arena consisted of a 40 × 40 × 40 cm^3^ opaque white foamboard Plexiglas box with a screen embedded in the ceiling. Throughout the experimental sessions, the monitor displayed a dim gray background (40 lux), on which a dark expanding disk was presented as the looming stimulus during the *Exposure* phase (Fig. 1A). Each stimulus consisted of ten expansions of a dark disk projected onto the gray background of the monitor. A single disc expansion consisted of two phases: 1) the disk expanded over a duration of 0.5 seconds, 2) the disk then remained static at its maximum expansion for 0.5 seconds before disappearing. An interval of 0.5-second was left before the next disk expansion; thus, each stimulus presentation lasted 10 seconds. Each mouse was exposed once, to one unique condition: 0, 6.6, 13.1, 27.5 or 36.5 % Michelson contrast.

Mice were dark-adapted for 30 min and then placed in the LTTA arena for 4 minutes (*Pre-Exposure*, Fig. 1A). A single contrast of looming stimulus of contrasts ranging from 0-36.5% Michelson contrast and then recorded in the arena for 1 minute following the *Exposure* phase. After a retention period of 2 days in the home cage, 30-minutes dark-adapted animals were then placed in the same arena for 4 minutes with no stimulus triggered (*Test* phase, Fig. 1A) and behavior was recorded using two cameras (C920x HD, Logitech and ELP Wide Angle Infrared Webcam).

For chemogenetic experiments, 4 weeks after viral infections (see below), mice were dark adapted and 30 min before the *Pre-Exposure* and *Test* phases of the LTTA paradigm, animals were intraperitoneally (i.p.) injected with saline (Vehicle, 0.9 % NaCl) or Clonazepine N-Oxide (1 mg/kg) (CNO, Tocris Bioscience Cat# 6329). After i.p. injections behavioral assay was performed as described above.

### Behavior analysis

For behavior analysis, we used eZtrack automated tracking software (Pennington et al., 2019) to analyze IR videos and quantify freezing behavior during *Exposure* phase of the LTTA paradigm at different looming stimuli contrasts. Freezing behavior was computed when mice were tracked immobile for at least 1 second. To quantify avoidance behavior, we used eZtrack software to analyze IR and regular light videos and quantify the time spent in the center of the arena during the *Test* phase (TZ_test_) of the LTTA paradigm. This parameter was normalized to the time spent in the center of the arena in the *Pre-Exposure* phase for each mouse (TZ_pre-exp_).

### Estrous cycle tracking

Estrous cycle tracking in female mice was performed after *Exposure* and *Test* phases of the LTTA paradigm (Fig. 5C-D). Estrous cycle phases were determined using vaginal lavages and histological analysis of the smears as previously reported (McLean et al., 2012). Briefly, for vaginal lavage female mice were immobilized and ∼25-50 μl of PBS 1x was expelled at the opening of vaginal canal using a latex bulb attached with a filtered 200 μl tip. The fluid was then placed on a charged glass slide and the smears were dried at a hot plate. Smears were then stained with Cristal Violet 0.1% (Sigma, C0775) and imaged on a Leica SP5 microscope in bright field mode. Vaginal cytology was performed, and the relative ratio of cell types observed in smears were used to identify the stage of the estrous cycle of mice on the LTTA phase of sample collection. During *Proestrus*, cells are almost exclusively nucleated epithelial cells (Fig. 5D, black arrowhead). In *Estrus* phase, cells are predominantly cornified squamous epithelial cells (Fig. 5D, black arrow). During *Metestrus*, small darkly stained leukocytes predominate (Fig. 5D, red arrowhead) with fragments of cornified squamous epithelial cells. During *Diestrus*, predominating leukocytes (Fig. 5D) and rare epithelial and cornified cells may still be present. For behavior analysis, mice were grouped into two categories: sexually receptive (SR -*Proestrus* and *Estrus*) and non-sexually receptive (NR -*Metestrus* and *Diestrus*) mice.

### Viral infection

Mice were anesthetized with isoflurane (Kent Scientific VetFlo system) and then unilaterally or bilaterally injected with 150 nl of AAVs (see Table S1) in the perihabenular nucleus (PHb) (AP:-1.60, ML:±0.55, DV:2.65 mm), ventral lateral geniculated nucleus (vLGN) (AP:-2.15, ML:±2.65, DV:3.60 mm), nucleus accumbens (NAc) (AP:1.42, ML:±0.80, DV:4.00 mm) or ventromedial prefrontal cortex (vmPFC) (AP:1.94, ML:±0.40, DV:2.82 mm) using a stereotaxic injector (Neurostar) controlled by the software Stereodrive at an injection rate of 30 nl/min. For intravitreal injections, mice were anesthetized by intraperitoneal (IP) injection of Avertin (2,2,2-Tribromoethanol) and a 30-gauge needle was used to open a hole in the *ora serrata*. Each eye was intravitreally injected with 1 μL of AAVs (see Table S1) using a custom Hamilton syringe (Borghuis Instruments) with a 33-gauge needle (Hamilton). After surgery procedures, mice were subcutaneously injected with 2 mg/kg of meloxicam (Covetrus). Efficiency of the viral injections were tested *postmortem*.

### Immunohistochemical procedures

For immunological studies in brain sections, mice were anesthetized by IP injection of Avertin and intracardially perfused with saline solution (PBS) followed by PFA 4% in PBS. Brians were dissected and post-fixated in PFA 4% in PBS overnight at 4 °C. For cryoprotection, brains were incubated in sucrose 30% for 3 days, embedded in OCT and then sectioned at 40 µm using a Leica CM1950 cryostat. For immunolabeling studies in retinas, animals were anesthetized by IP injection of Avertin and sacrificed by cervical dislocation. Eyecups were enucleated and retinas were dissected and then fixed in PFA 4% in PBS for 30-60 minutes at RT. Brain slices and retinas were washed and then blocked at 4 °C overnight in 6% normal goat serum in 0.3% Triton PBS prior to incubating in primary antibody solution for 2-3 nights at 4 °C. Then, tissue was washed and incubated in secondary antibody solution for 2 hours at RT (Table S2). The tissue was finally mounted using Fluoromount (Sigma) and imaged using a Leica SP5 confocal microscope. Primary and secondary antibody solutions were made in 3% normal goat serum in 0.3% Triton PBS. For PHb-ipRGC mCherry expression analysis, high magnification images (pixel size of 0.36 µm) with a z-stack size of 1 µm were taken. For viral expression in brain regions, high magnification images (pixel size of 0.72 µm) with a z-stack size of 2 µm were taken. All images were processed and analyzed using ImageJ.

### Fiber photometry

Mice were anesthetized with isoflurane (Kent Scientific VetFlo system) and then unilaterally injected with 150 nl of AAV1-CAG-GCaMP6s-WPRE-SV40 (∼8 × 10^12^ GC/mL, see Table S1) in the PHb using a stereotaxic injector (Neurostar) controlled by the software Stereodrive at an injection rate of 30 nl/min. A FP cannula (MFC_200/230-0.48_3mm_MF1.25_FLT, DORIC) was then implanted 150 µm above the site of injection. After surgery procedures, mice were subcutaneously injected with 2 mg/kg of meloxicam (Covetrus). Efficiency of the brain injection and fiber implantation were tested *postmortem*.

Behavioral assay and FP recordings were made 4 weeks post-surgery. For that, dark-adapted mice were exposed to the LTTA paradigm as described above. For FP recording, an optic fiber (MFP_400/430/1100-0.57_1m_FC-ZF1.25(F)_LAF, DORIC) was attached to the head of the animals and then were placed to the LTTA arena. FP signal was collected using an assisted electrical rotary system (AERJ_24_FMC, DORIC), Amplifier (Doric Fluorescence Detector, DORIC), LED driver (Laser Diode Fiber Light Source, DORIC) and processor (RZ5P Base Processor, Tucker-Davis Technologies). FP data was extracted using pMAT software (Bruno et al., 2021). All FP data processing and behavior correlation analyses were made using eZtrack and custom phyton scripts.

### Statistical comparisons

All graphs and statistical analyses were performed using Graph Pad Prism 10.1.2 (RRID: SCR_002798). Normal distribution of data was tested using Shapiro-Wilk test. Outliers were identified using ROUT (Q=1) method and excluded from the data analysis. Student’s t-or Mann Whitney U tests were used to compare two experimental groups. For multiple statistical comparisons, we performed one-way ANOVA followed by Tukey’s and Dunn’s tests. Significance was concluded when P < 0.05.

## Supplemental information

**Table S1.**
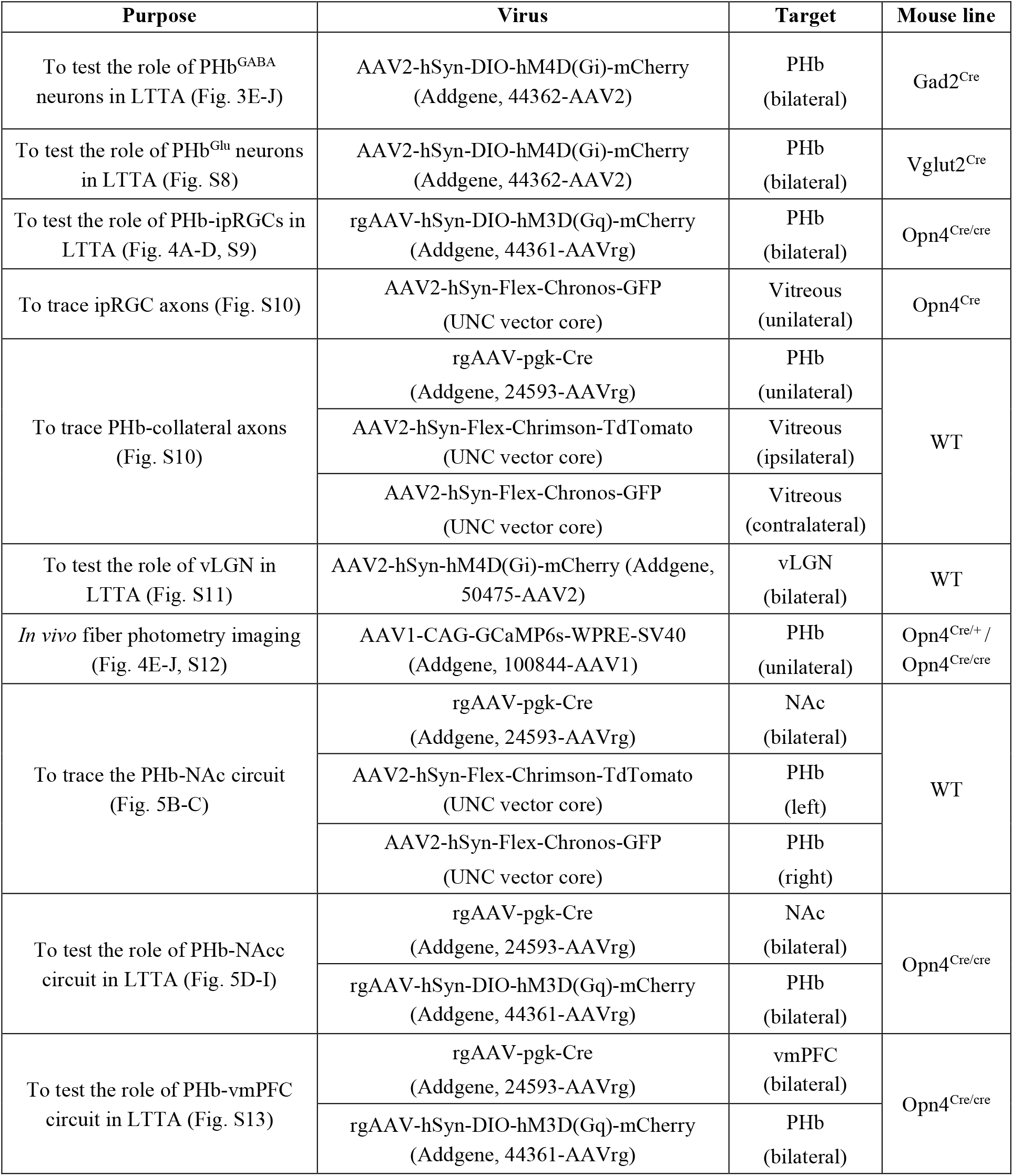
List of intersectional virus strategies used in this study.

**Table S2.** List of antibodies used in this study. The dilution of all secondary antibodies was 1:500.

| **Purpose** | **Primary Ab** | **Dilution** | **Secondary Ab** |
| --- | --- | --- | --- |
| To find candidate brain regions that drive LTTA (Fig. 3B-C, S7) | Rabbit anti-cFos<br>(Cell signaling, 2250) | 1:500 | Donkey anti-rabbit Alexa 488<br>(Thermofisher, A-21206) |
| To characterize viral expression carrying mCherry or TdTomato reporters (Fig. 3F, 4B, 5C, 5F, S8, S11, S13) | Rabbit dsRed<br>(Takara, 632496) | 1:500 | Donkey anti-rabbit Alexa 594<br>(ThermoFisher, A-21207) |
| To characterize viral expression carrying GFP reporter (Fig. 5C, S10, S12) | Chicken anti-GFP<br>(Abcam, ab13970) | 1:500 | Donkey anti-chicken Alexa 488<br>(Jackson IRL, 703-545-155) |

## Notes

### Competing Interest Statement

The authors have declared no competing interest.

